# Experiencing surprise: the temporal dynamics of its impact on memory

**DOI:** 10.1101/2020.12.15.422817

**Authors:** Darya Frank, Alex Kafkas, Daniela Montaldi

## Abstract

To efficiently process information, the brain shifts between encoding and retrieval states, prioritising bottom-up or top-down processing accordingly. Expectation violation before or during learning has been shown to trigger an adaptive encoding mechanism, resulting in better memory for unexpected events. Using fMRI we explored (1) whether this encoding mechanism is also triggered during retrieval, and if so, (2) what the temporal dynamics of its mnemonic consequences are. Male and female participants studied object images, then, with new objects, they learned a contingency between a cue and a semantic category. Rule-abiding (expected) and violating (unexpected) targets and similar foils were used at test. We found interactions between previous and current similar events’ expectation, such that when an expected event followed a similar but unexpected event, its performance was boosted, underpinned by activation in the hippocampus, midbrain, and occipital cortex. In contrast, a sequence of two unexpected similar events also triggered occipital engagement, however, this did not enhance memory performance. Taken together, our findings suggest that when the goal is to retrieve, encountering surprising events engages an encoding mechanism, supported by bottom-up processing, that may enhance memory for future related events.

**Significance statement:** Optimising the balance between new learning and the retrieval of existing knowledge is an ongoing process, at the core of human cognition. Previous research into memory encoding suggests experiencing surprise leads to the prioritisation of the leaning of new memories, forming an adaptive encoding mechanism. We examined whether this mechanism is also engaged when the current goal is to retrieve information. Our results demonstrate that an expectation-driven shift towards an encoding state, supported by enhanced perceptual processing, is beneficial for the correct identification of subsequent expected similar events. These findings have important implications for our understanding of the temporal dynamics of the adaptive encoding of information into memory.

## Introduction

To efficiently process inputs, the brain shifts between top-down and bottom-up streams, balancing processing of sensory inputs versus utilisation of stored representations. This is reflected by the existence of distinct encoding and retrieval states (Buzsáki, 2002). Bottom-up processing supports the hippocampal encoding state, prioritising transformation of perceptual inputs into memories, while top-down processing supports a retrieval state by facilitating access to stored information. The specific mechanisms that govern encoding-retrieval shifts, which may or may not function competitively, are not well understood, but will be driven, at least in part, by the factors that induce each memory state. Whilst there is ample evidence that an adaptive encoding state is triggered upon experiencing discrepancies between expected and encountered events, it remains unclear whether such an implicit learning mechanism might also be engaged when the dominant goal is to retrieve, rather than encode. Using behavioural (Experiment 1) and fMRI (Experiment 2) data, we examine whether surprise produced by expectation violation at retrieval, results in a shift towards an encoding state, accompanied by increased perceptual processing; and whether this occurs at the expense of retrieval. We also investigate the extended consequences of this expectation violation, and the potential encoding-retrieval iterative shifts it might generate, by examining trial-by-trial recognition of subsequent similar events.

Surprise, produced by expectation violation, has been shown to engage hippocampal encoding (Axmacher et al., 2010; Kafkas and Montaldi, 2015; Long et al., 2016; Frank et al., 2020), which together with the midbrain dopaminergic system (Lisman and Grace, 2005; Shohamy and Wagner, 2008; Kafkas and Montaldi, 2018a) and enhanced perceptual processing (Stoppel et al., 2009; Hawco and Lepage, 2014; Kafkas and Montaldi, 2014, 2015), supports adaptive memory formation. Evidence for this adaptive mechanism comes mostly from paradigms in which expectation violation takes place before or during learning (Li et al., 2003; Garrido et al., 2015; Long et al., 2016; Greve et al., 2017; Kafkas, 2021). Therefore, any additional resources diverted towards encoding in these scenarios is likely to boost later memory performance (e.g. attention effects see Aly and Turk-Browne, 2017). However, to demonstrate the ubiquity of an adaptive encoding mechanism, it is critical to provide evidence of a shift towards encoding also during retrieval. This is likely to shed light on the effects of the rapid and implicit real-life interplay between these states. Under these circumstances, engaging encoding still serves an adaptive purpose, but it might not be ‘beneficial’ in real-time.

Given the malleability of memories triggered by surprise (Kim et al., 2014), it becomes important to consider what the mnemonic consequences of a shift towards encoding may be. Should expectation violation trigger an encoding state, at the expense of a memory search, it might modulate processing of the unexpected event. This is consistent with the view that the hippocampus continuously shifts between encoding and retrieval states (Buzsáki, 2002; Hasselmo et al., 2002). This system can be likened to a pendulum swinging rhythmically between the two memory states, optimising information processing. Should situational factors disrupt the on-going swing between states, favouring one over another, the result can be a change in memory efficiency. Previous research has shown that an explicit strategy cue fosters a hippocampal trade-off between encoding and retrieval of information (Richter et al., 2016).

We examined whether expectation violation at retrieval spontaneously engages an encoding state, and if so, whether this response is dependent on the perceptual similarity between inputs. We also considered the sustained mnemonic consequences of such iterative shifts on current and subsequent recognition decisions. Expectation was manipulated implicitly and independently of the requirements of the retrieval task, allowing us to address two important questions; first whether expectation violation engages a bottom-up encoding mechanism, even when it is not goal-relevant. Second, what are the temporal dynamics of the mnemonic consequences of expectation status, for an event and subsequent similar events. We reasoned that heightened expectation-driven encoding might modulate current retrieval processes, but aid accuracy of recognition of subsequent similar events.

## Methods

### Experiment 1

#### Participants

30 participants (4 males) between the ages 18-22 (M = 19, SD = 1.04) gave informed consent and took part in the experiment. The sample size was selected based on previous studies utilising a similar design and analysis approach (Kafkas and Montaldi, 2018b; Frank et al., 2020). Participants had normal or corrected-to-normal vision and no history of neurological or psychiatric disorders. All procedures were approved by the University of Manchester Research Ethics Committee. Four participants were excluded from any further analysis due to failure to learn the cue-outcome association (3) or recognition performance below chance (1). Data from 26 participants are reported.

#### Materials

78 images of natural (39) and man-made (39) objects were selected from the Similar Objects-Lures Image Database (SOLID; Frank et al., 2019). These images were used as the target objects, presented during encoding. Using the dissimilarity index from SOLID, three foils of decreasing levels of similarity (F1 – most similar, F2 – mid-level F3 - lowest similarity) were selected for each target image. Additionally, similarity was parametrically manipulated by keeping the average distance between the levels constant (average dissimilarity 2100 DI; see Figure 1 for examples and Frank et al., 2019 for further explanation on DI values). These levels of similarity were chosen based on our previous results (Frank et al., 2020), where we observed a quadratic pattern of expectation effects on inputs similarity, when expectation was manipulated at encoding, with respect to memory performance. Therefore, 78 object sets (one target and three foils; total 312 stimuli) were utilised.

**Figure 1.**
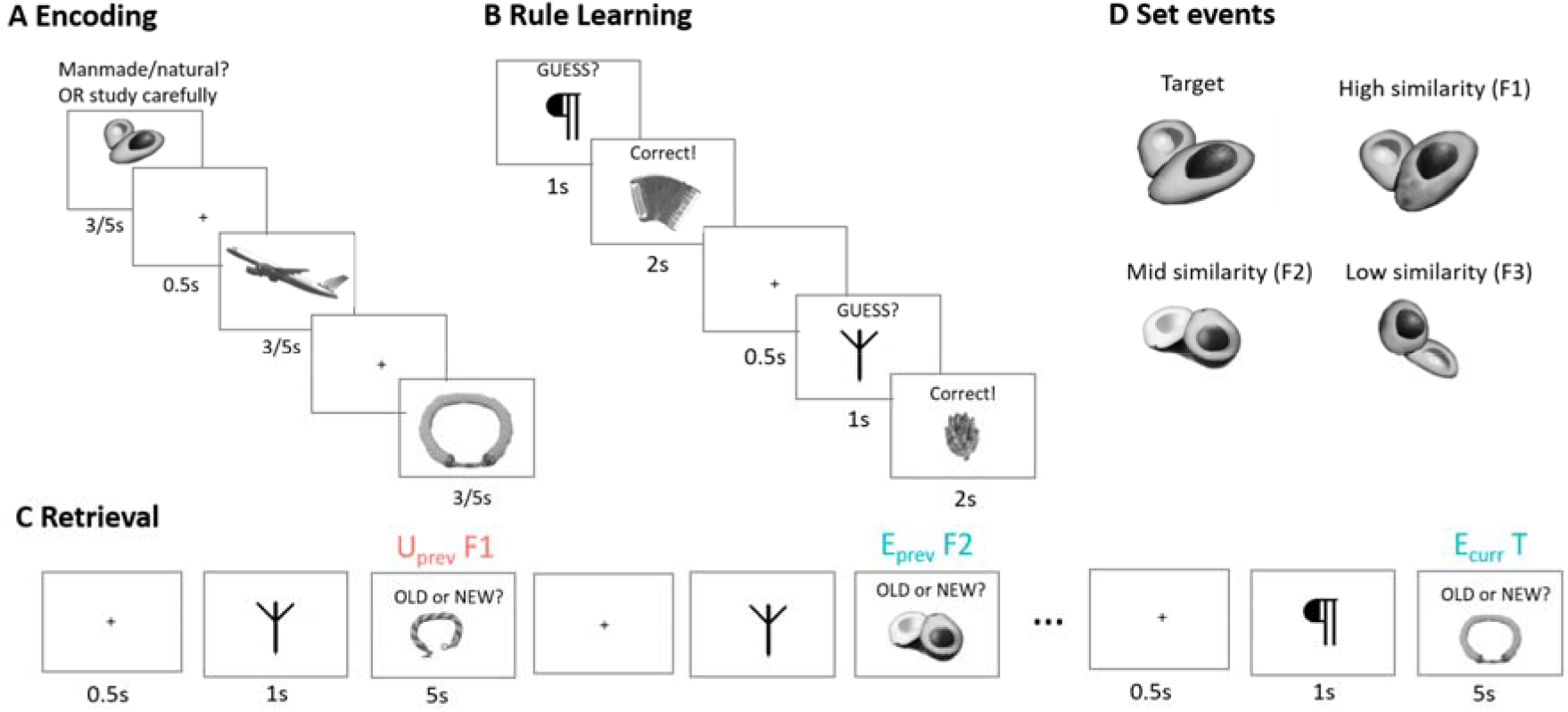
Experimental design. **A)** During the first round of **encoding**, participants responded whether the object was man-made or natural. In the second encoding round, participants are asked to study the object carefully. In Experiment 2, each object appeared three times consecutively; in addition to the two presentations from Experiment 1, participants were asked to respond whether the object was more likely to be found indoors or outdoors. The order of man-made/natural and in/outdoor questions was randomised, the third presentation was always ‘study carefully’. **B)** In the **rule-learning** task participants learned a contingency between a cue and an object’s category, man-made or natural. Participants had 1 second to make a decision in Experiment 1, and 3 seconds in Experiment 2. See Extended Data Figure 1-1 and 1-2 for rule-learning performance. **C)** In the **retrieval task**, the same cues were presented before each set event, old objects (targets) and new similar foils (F1, F2, F3 in Experiment 1; F1 and F2 in Experiment 2). On 30% of these trials there was a mismatch between the cue and the object’s category, these are unexpected trials (e.g., a cue for a natural object is followed by a man-made object, marked in red). Participants were instructed to indicate whether the object is old or new. U_*prev*_*F1, EcurrT* = Unexpected F1 presented before an expected target, from the same set. **D) Example of range of perceptual similarity** within set events.

#### Experimental design

The experiment was controlled using PsychoPy version 1.82 (Peirce, 2007) and consisted of four parts (similar to design used in Kafkas and Montaldi, 2018b). First, at encoding, participants were presented with the target objects twice to ensure sufficient exposure. During the first presentation, the object was shown on the screen for 3 seconds, and participants were asked to decide whether the object was man-made or natural; pressing the left arrow key for natural, and the right arrow key for man-made. In the second presentation participants were asked to pay close attention to the images (shown for 5s each) and were informed that they would be asked to distinguish between the presented (‘old’) object and similar (‘new’) objects later; they did not have to make any response during the second encoding presentation. The order of image presentation was randomised across participants. The next phase involved a 5-minute filled delay task, during which participants solved arithmetic problems.

In the third phase, a rule-learning task was used to allow manipulation of expectation later at retrieval. Participants were asked to learn the contingency between a symbol (acting as a cue) and an object’s category, man-made or natural. Four symbols were used in total, two for each category, and these were counterbalanced between participants. Each cue was presented 14 times. Trials began with a 500ms fixation point, followed by the cue, presented for 1 second. During this time participants were instructed to guess the following object’s category by using the same keys as in the encoding phase. Feedback and a new object (not tested) were then presented together for 2 seconds. This task established participants’ expectation regarding the cue-object sequences, and critically this was manipulated at retrieval. To ensure expectations were set, only data from participants who reached criterion (above 50% accuracy in the first half, and 75% accuracy in the second half of the task) were analysed (See Supplementary Figure S.1 for accuracy and RT in the rule-learning task).

The final phase was an old/new recognition task, during which all set events (targets, F1, F2 and F3) were presented. The experimenter informed participants that they will be presented with old (target) objects, and similar new ones (foils). Each retrieval trial began with a fixation point (500ms), followed by a cue (1 second) and an object (up to 5 seconds). Participants were told to focus on the object and press ‘old’ if they thought it was exactly the same as the target they had previously seen. Participants were instructed to press ‘new’ if they noticed anything different in the object. Following 12 practice trials, the main task began. The key manipulation at retrieval was the validity of the cue. One-third of the cues were misleading, making the object to-be-recognised unexpected. Valid trials (e.g. cue natural followed by a natural object) were marked as expected. As an old/new recognition task was used, the four old/new decisions per object set were independent of each other, critically meaning that an ‘old’ response could be given for multiple set events (i.e. for target and one or more foils). After completing the experiment participants were debriefed regarding the expectation manipulation at retrieval. Only two participants in Experiment 1 and five participants in Experiment 2 reported noticing the mismatch between cue and object during the task.

#### Statistical analysis

To assess expectation-modulated dynamic encoding, we collated object sets (apples, scissors etc.). In each set, there were four set events (target, F1 - most similar, F2 – mid similarity, and F3 – least similar), each with a randomly assigned (a) expectation condition, (b) presentation position within the set, and (c) order of presentation at retrieval (1-312 trials). We ran a mixed-effects binary logistic regression model on these ungrouped data, using expectation status of the current and previous set events, and the presentation position at retrieval as a covariate. Models were computed using the lme4 package (Bates et al., 2015) in the R environment (R Development Core Team, 2008). The parameters of such models can be used to assess the probability of giving a correct response (‘old’ for targets, ‘new’ for foils) and also account for each participant’s unique intercept (response bias). To assess the slope of each predictor in the model (H_0_: β = 0), we used an omnibus χ^2^ Wald test (West et al., 2014), as implemented in the car package (Fox and Weisberg, 2018). Extraction and plotting of the effects reported below was conducted using the effects (Fox, 2003), emmeans (Searle et al., 1980) and ggplot2 (Wickham, 2009) packages. To examine the dynamic interaction between perceptual similarity and expectation status, for each set event we devised three models of interest, modelling each event separately as a function of the preceding event sequence. For example, targets preceded by F1 were modelled separately from targets preceded by F2. To eliminate any effects driven by memory strength differences, we only included in this model events for which the previous response was correct (i.e. for targets following F1 events, we only included targets that followed CR1). Similar results were observed when including all trials in the model (see GitHub repository for code to run analyses and generate figures). Each model thus included the current and previous set events’ expectation status, as well as the order of presentation at retrieval as a covariate. Code and data are available here: https://github.com/frdarya/DynamicExpectation.

#### Experiment 2

##### Participants

25 participants (8 males, ages 18-33, M = 25, SD = 4.2) gave informed consent and took part in the study. Participants had normal or corrected-to-normal vision and no history of neurological or psychiatric disorders. All procedures were approved by the University of Manchester Research Ethics Committee. One participant was excluded from all analyses due to failure to learn the cue-outcome contingency during the rule-learning task.

##### Experimental design

A similar paradigm and expectation manipulation was used in Experiment 2, with the following exceptions: in the encoding phase, each object was presented three times consecutively. Each object was on the screen for 2000ms, with a jittered fixation cross (250-750ms) between each presentation. During the first and second presentations participants were asked to make a semantic decision about the object, whether it is man-made or natural, and whether it is more likely to be found indoors or outdoors. The order of these questions was random. During the third presentation participants were always asked to study the object carefully focusing on the details. Following the third presentation there was another jittered fixation cross, for a longer period of time (800-1200ms), to create mini-blocks separating each object.

The rule-learning task was identical to that used in Experiment 1, except for a longer response presentation time of the cue (3s instead of 1s) and a jittered fixation cross (250-750ms) between each trial (see Supplementary Figure S.1 for behavioural and Figure S.2 for fMRI results from the rule learning task). Before the retrieval task, participants solved arithmetic problems for two minutes (not scanned). At retrieval, we used two levels of foil similarity (F1 and F2) in each set, instead of three. In the interest of optimising time-in-scanner, F3 objects were not used, as they did not yield any effects of interest in Experiment 1. Each retrieval trial started with a jittered fixation cross (250-750ms), followed by a presentation of the cue for 1000ms and then the set event (target, F1 or F2) for 3000ms. In all scanned tasks, we used implicit baselines (fixation crosses for 3500ms in encoding and rule-learning tasks, 4500ms in retrieval) in 30% of trials.

##### Behavioural statistical analysis

Following the analysis and results from Experiment 1, we collapsed targets and F1 events and modelled the probability of making a correct decision (hits and correct rejections) based on the current set event’s expectation status and the previous set event’s expectation status. As in Experiment 1, to eliminate the memory strength confound, only correct responses were included (see GitHub repository for similar results when including all trials, and for separate analyses of targets and F1). Correct rejections of F2 foils were also modelled as a function of previous set events, as was done for Experiment 1.

##### fMRI acquisition and statistical analysis

MR scanning was carried out on a 3T MRI scanner (Philips, Achieva). To minimise movement during the scan, foam wedges and soft pads were used to stabilise the participant’s head. First, T1-weighted images (matrix size: 256×256, 160 slices, voxel size 1mm isotropic) were collected while participants rested in the scanner. A gradient echo-planar imaging (EPI) sequence was used to collect T2* images for the BOLD signal. 40 slices parallel to the AC-PC line, covering the whole brain (matrix size 80 × 80, voxel size 3 × 3 × 3.5mm^3^), were obtained for each volume (TR = 2.5s, TE = 35ms). Participants performed three tasks in the scanner (encoding 313 volumes; rule-learning 143 volumes; retrieval 534 volumes) and a distractor task, which was not scanned.

fMRI data were pre-processed and analysed using SPM12 (Statistical Parameter Mapping, Wellcome Centre for Human Neuroimaging, University College London https://www.fil.ion.ucl.ac.uk/spm/software/spm12/). Images were realigned to the mean image using a six-parameter rigid body transformation, resliced using sinc interpolation and slice-time corrected to the middle slice. T1 anatomical images were co-registered to the corresponding mean EPI image. Spatial normalisation to the Montreal Neurological Institute (MNI) template was carried out using the DARTEL toolbox implemented in SPM12 (Ashburner, 2007). An isotropic 8mm FWHM Gaussian kernel was used for smoothing the normalised EPI data for univariate analyses. To remove low-frequency noise the data was high-pass filtered using a cut-off of 128s. Two a-priori regions of interest (ROIs), the bilateral hippocampus, and a midbrain ROI including only the substantia nigra (SN) and ventral tegmentum area (VTA) were used. The hippocampus mask was taken from the Harvard-Oxford anatomical atlas (threshold at 25% probability; Desikan et al., 2006). The midbrain mask was taken from the probabilistic atlas of the midbrain (Murty et al., 2014).

Each participant’s functional data from the retrieval session was analysed using the general linear model (GLM) framework within an event-related design modelling the canonical hemodynamic response function. The six motion parameters produced at realigmnet for each session were used as nuisance regressors. To minimise residual motion artefacts the ArtRepair toolbox (http://cibsr.stanford.edu/tools/human-brain-project/artrepair-software.html) was used to produce additional nuisance regressors for each participant. The time series were high-pass filtered to remove low frequency noise (128s cut-off). Given our a priori hypothesis for the ROIs introduced above (bilateral hippocampus and SN/VTA), a small volume correction (SVC) approach was adopted for these regions, corrected for family-wise error (FWE) for the ROI volume (initial cluster-forming threshold of p < 0.001). For more exploratory whole-brain analyses, a non-parametric permutation-based (n = 5000) approach was used to identify significant clusters using the SnPM toolbox (https://warwick.ac.uk/fac/sci/statistics/staff/academic-research/nichols/software/snpm/). An initial cluster-forming threshold of p < 0.005 was used, and clusters significant at a non-parametric p < 0.05 are reported. Unthresholded group-level T-maps from SPM and SnPM are available here: https://neurovault.org/collections/TTDMPHLE/.

To test the behavioural effect of an interaction between a previous event’s contextual expectation and the current event’s contextual expectation, we collapsed targets and F1 and classified them based on presentation order (which came first) and their expectation status. F2 events were modelled as a separate condition. In this analysis, we compare the current item between current and previous expectation status (e.g. **E**_prev_**U**curr > **E**_prev_**E**curr). Given our experimental design, there are four parameters whose interactions could be further explored: set event (target, F1, F2), contextual expectation (expected or unexpected), memory response given (correct or incorrect) and presentation order within the set (first, second or last). This results in 36 conditions, however, these could not be modelled together due to insufficient number of trials per bin (n < 7) for the majority of participants. Therefore, we devised three separate models; in the first model, we examined the main effect of expectation, irrespective of set event or recognition decision. In the second model, we examined expectation x set event x successful memory interactions by collapsing trials across presentation order, and modelling only correct responses for each event (all incorrect responses were modelled as a separate regressor). In the third model, we explored the interaction of expectation, set event and presentation order, irrespective of recognition responses.

## Results

We conducted two experiments, Experiment 1 examined behavioural responses, and Experiment 2 employed a similar paradigm whilst fMRI data was collected. In both Experiments, our behavioural task included three stages (see Figure 1 for experimental design): following the encoding of object images, participants performed a rule-learning task in which they learned a contingency between a cue and the object’s category (man-made or natural). Then, at retrieval, the same cues were presented, followed by an old (target) or new (foil) object. Foils were parametrically manipulated similar objects, F1, F2, F3 in a decreasing order of similarity to the target. Participants were asked to make an old/new recognition decision. In one-third of the retrieval trials, the pre-established cue and object category contingency was violated. To assess expectation-modulated encoding, and its sensitivity to perceptual similarity, we used mixed-effects logistic regression to model the response (correct/incorrect recognition) to each set event (targets and similar foils) as a function of the preceding event from that set (e.g. if first an F1 is presented, and later on a target, these would be noted F1_prev_ Target_curr_). Furthermore, each set event was associated with an expectation status determined by the preceding cue (rule-abiding events were marked as expected rule-violating as unexpected). Each model also included an interaction between current and previous set events’ expectation status to examine dynamic changes (e.g. how a previous unexpected foil affects recognition of a current expected target; see Figure 1c, and Methods for full details).

### Experiment 1

#### Memory for targets is modulated by preceding unexpected similar set events

For targets following F1 set events, we found significant main effects of the F1 expectation (β = 0.659, Χ^2^(1) = 4.63, p = 0.031), as well as a marginal interaction between the target’s expectation and the previous F1’s expectation status (β = -0.939, Χ^2^(1) = 3.65, p = 0.056; Figure 2a). Subsequent contrast tests revealed that expected targets were more likely to be remembered following an unexpected F1, compared to an expected F1 set event (z = 2.15, p = 0.031). Furthermore, when the previous F1 set event was unexpected, the subsequent expected targets were more likely to be correctly remembered, compared to unexpected targets (z = 2.37, p = 0.017). When examining targets that followed F2 and F3 set events, we did not observe any significant predictors (all p’s > 0.127).

**Figure 2.**
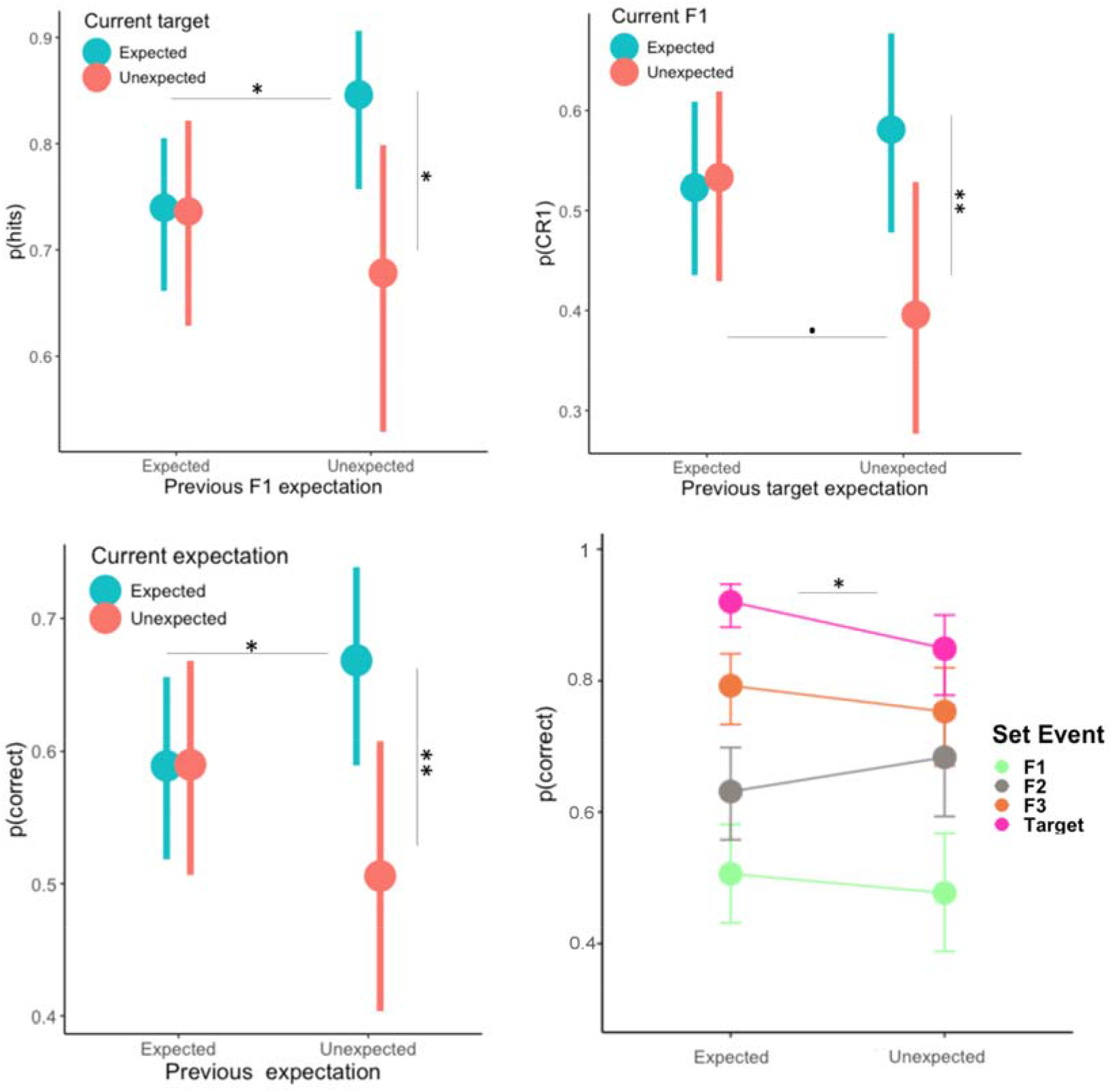
Experiment 1 Results. **A) Predicting hits**. More hits were observed for expected targets following unexpected F1 foils, compared to unexpected targets following unexpected F1, and compared to expected targets following expected F1. **B) Predicting CR1**. More CR1 for expected F1 following unexpected targets, compared to unexpected F1 following unexpected targets, and a marginal effect of poorer memory (less CR1) for a sequence of unexpected events. **C) Collapsed hits and CR1**. Similar results showing an interaction between the previous and current set events’ expectation status, with U_prev_E_curr_ events showing the best memory performance, compared to U_prev_U_curr_ and E_prev_E_curr_. **D) First set events**. More hits for expected compared to unexpected targets presented first in the event sequence. No other significant effects, although F1 events follow a similar direction. Unless otherwise stated, all error bars represent 95% confidence intervals. • p = 0.0507, * p < 0.05, ** p < 0.01.

#### Correct rejection of similar foils is modulated by the expectation status of preceding targets

For F1 events following targets, we found an interaction between the two events’ expectation status (β = -0.79, Χ^2^(1) = 5.29, p = 0.021; Figure 2B), with contrasts showing more correct rejections of F1 events (CR1), for expected F1 than unexpected F1 events following unexpected targets (z = 2.65, p = 0.008) and marginally less correct rejections for unexpected F1 when the previous target was unexpected, compared to when it was expected (z = 1.95, p = 0.0507; Figure 2 middle panel). All other effects were not significant (all p’s > 0.255). When examining F1 events that followed F2 and F3 set events, we did not observe any significant predictors (all p’s > 0.274).

#### Expectation sequence modulation decreases as perceptual similarity decreases

F2 events were not affected by previous targets (all p’s > 0.171), F1s (all p’s > 0.197), or F3s (all p’s > 0.174) from the same set. Similarlary, responses to F3 events were not modulated by any preceding set events (targets: all p’s > 0.484, F1s: all p’s > 0.148, F2s: all p’s > 0.321).

#### Targets and their most similar foils show analogous expectation sequence effects

Given the similar effects observed for targets and F1 events independently, we collapsed the two, to examine whether these effects are complementary (i.e., whether there is an interaction between current and previous expectation status; Figure 2C). Although hits and correct rejections are not necessarily products of the same mnemonic process, in this paradigm they provide an opportunity to examine how perceptual load (in the form of similarity) interacts with dynamic expectation modulations. The mnemonic comparison between the current set event (target or F1) and its previous set event (F1 or target, respectively), forms the highest load, or interference, in relation to the encoded object, as the participant makes a recognition decision. Therefore, if perceptual processes are engaged, upon encountering an unexpected event, the effects observed for each set event individually should replicate. We found a significant main effect of previous event’s expectation status (β = 0.341, Χ^2^(1) = 4.9, p = 0.027), as well as a significant interaction between expectation status of the current and previous events (β = -0.682, Χ^2^(1) = 6.5, p = 0.01). Subsequent contrast tests revealed that when the previous set event was unexpected, more correct responses were observed for expected compared to unexpected events (U_prev_E_curr_ > U_prev_U_curr_; z = 3.1, p = 0.002). U_prev_U_curr_ was numerically smaller than E_prev_U_curr_, but this effect did not reach statistical significance (p = 0.116). Additionally, there were more correct responses for current expected events following unexpected than expected events (E_prev_E_curr_ < U_prev_E_curr_; z = -2.22, p = 0.027).

Taken together, these results suggest expectation violation shifts processing away from retrieval and towards encoding, and that the temporal dynamics of the mnemonic consequences of this shift are reflected in the memory for the subsequent set event (effects of first set events depicted in Figure 2D), as a function of perceptual similarity. When subsequent events are unexpected (U_prev_U_curr_), we observed poor accuracy for F1 foils, and to a lesser extent for targets. On the other hand, when an unexpected event is followed by an expected event (U_prev_E_curr_) a boost in performance was observed for the current event, driven mainly by responses to targets. To examine whether these effects engage the circuit involved in adaptive memory formation (including the hippocampus and midbrain (Shohamy and Adcock, 2010; Kafkas and Montaldi, 2018a), as well as to test the hypothesis that expectation violation engages an encoding mechanism, supported by the bottom-up information stream (ventral visual pathway), in Experiment 2 a new set of participants performed a similar task while fMRI data was acquired (see Methods for minor task adjustments).

### Experiment 2

#### Behavioural Results

Replicating the effects observed in Experiment 1, we found a main effect of the previous set event’s expectation status (β = 0.309, Χ^2^(1) = 4.13, p = 0.042), as well as an interaction between the current and previous events’ expectation status (β = -0.59, Χ^2^(1) = 4.47, p = 0.034; Figure 3A). Subsequent contrast tests revealed better memory performance for expected events following unexpected ones, compared to those following expected events (U_prev_E_curr_ > E_prev_E_curr_; z = 2.55, p = 0.011). For set events following unexpected ones, better memory was also found for expected compared to unexpected events (U_prev_E_curr_ > U_prev_U_curr_; z = 2.68 p = 0.007). Next, we examined CR2 as a function of the previously seen targets and F1 events, and their expectation status. For F2 following targets, a main effect of the target’s expectation status was observed (β = 0.473, Χ^2^(1) = 4.4, p = 0.036), with more CR2 following unexpected targets. All other effects were not significant (all p’s > 0.169). Correct responses to F2 events were not affected by preceding F1 events (all p’s > 0.263).

**Figure 3.**
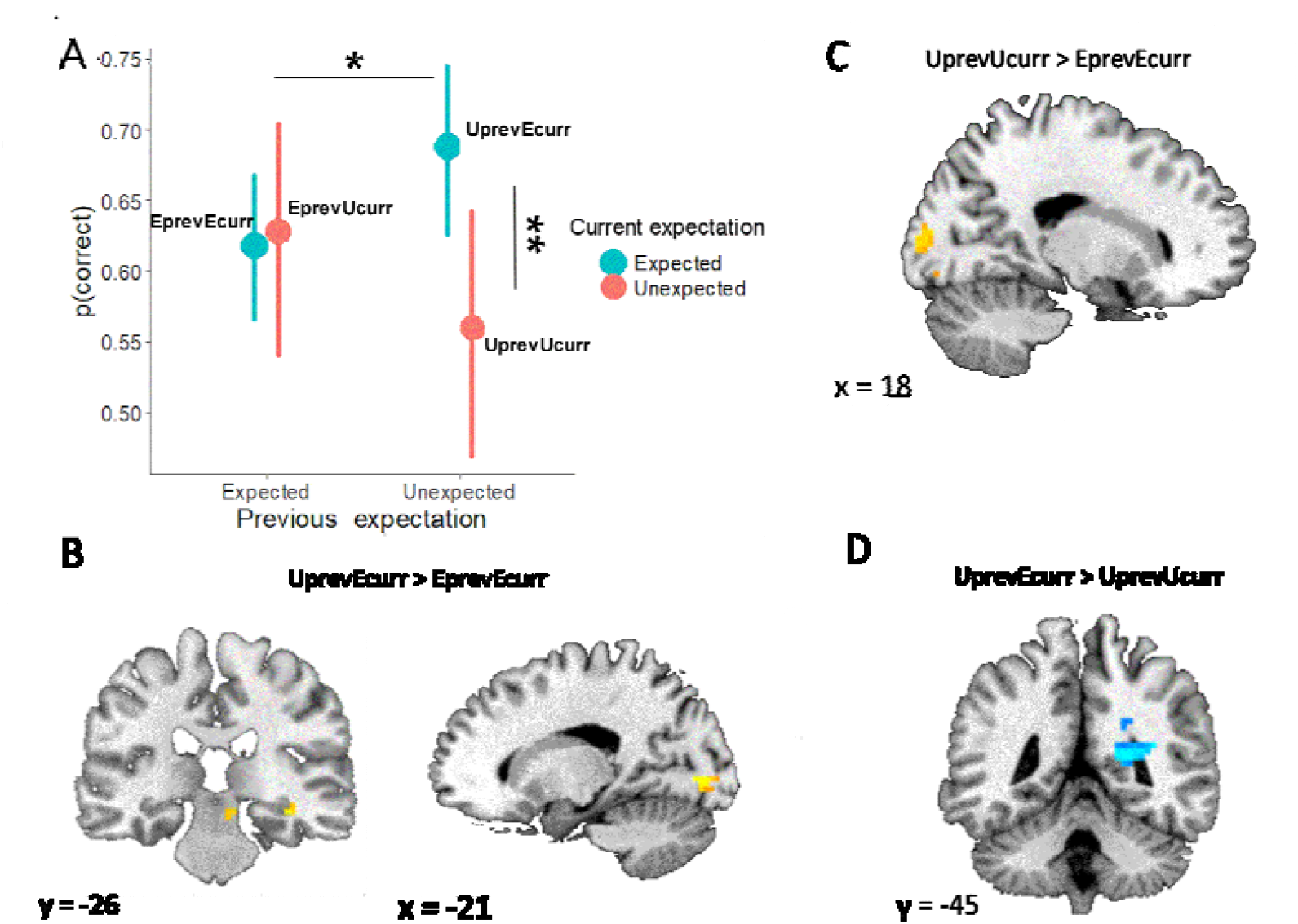
Behavioural and neural expectation interactions for targets and F1 foils. **A) Behavioural results**. Replicating the results from Experiment 1, a current by previous expectation status interaction was observed, with U_prev_E_curr_ showing a boost in memory performance. **B) UprevEcurr > EprevEcurr contrast**. Increased activation in the right hippocampus, SN/VTA and left inferior occipital cortex. **C) Unexpected > Expected interactions**. Increased activation in right occipital cortex (BA 18) was observed for U_prev_U_curr_ > E_prev_E_curr_, despite poor memory performance for U_prev_U_curr_ events. **D) Expected > Unexpected interactions**. Increased activation in right retrosplenial cortex/precuneus for U_prev_E_curr_ > U_prev_U_curr_.

#### fMRI results

##### Expectation sequence interactions engage hippocampal, midbrain, and occipital regions to support subsequent mnemonic processing

We first examined the neural correlates of the behavioural contextual expectation interaction reported above. For current expected events that followed unexpected events, compared to those following an expected event (U_prev_E_curr_ > E_prev_E_curr_; see Figure 3B), we found increased activation in the right hippocampus (x = 36, y = -33, z = -12, k = 12, SVC p_FWE_ = 0.04), SN/VTA (x = 9, y = -24, z = -12, k = 11, SVC p_FWE_ = 0.039), and left inferior occipital gyrus (BA 18; x = -21, y = -81, z = -18, non-parametric p_cluster_ = 0.018). For current unexpected events following previous expected ones, compared to those which followed an unexpected event (E_prev_U_curr_ > U_prev_U_curr_), reflecting poorer performance, we also found increased activation in the right hippocampus (x = 24, y = - 33, z = -6, k = 10, SVC p_FWE_ = .045) and left parahippocampus (x = -33, y = -45, z = -6, non-parametric p_cluster_ = 0.049). Critically, in both contrasts the current set events had the same expectation status and differed only on the expectation status of the previous event.

Comparing current expected vs. unexpected events, following previously unexpected events (U_prev_E_curr_ > U_prev_U_curr_; Figure 3d) revealed activation in the right retrosplenial cortex/precuneus (x = 24, y = -45, z = 12, non-parametric p_cluster_ = 0.0213). The complementary contrast, following expected events, E_prev_E_curr_ > E_prev_U_curr_ did not reveal significant effects. For the U_prev_U_curr_ > E_prev_E_curr_ contrast in which unexpected events elicited more activations than expected ones, despite showing reduced memory performance (Figure 3C), we found increased activation in right occipital cortex (BA 18, x = 18, y = -93, z = 12, non-parametric p_cluster_ = 0.0318). The complementary contrast U_prev_U_curr_ > E_prev_U_curr_ did not reveal significant effects. Comparing the first (previous) event between conditions (first expected vs. first unexpected) did not reveal any significant effects.

##### Expectation status differentially engages encoding and retrieval-related regions

To explore whether expected and unexpected events, across event types, responses, and temporal positions elicited differential activations in a bottom-up (ventral visual stream) or reinstatement (retrieval network) manner, we also compared the two conditions (see Figure 4A). We found increased activity for unexpected > expected events in right occipital cortex (BA 19, x = 39, y = -75 z = 12 and BA 18, x = 39, y = -75, z = 12, non-parametric p_cluster_ = 0.015) and right fusiform gyrus (x = 27, y = -48, z = -18, non-parametric p_cluster_ = 0.0173). For expected > unexpected, we observed increased activation in the right inferior parietal lobe (angular gyrus; BA 39 x = 48, y = -48, z = 33, non-parametric p_cluster_ = 0.0206) and bilateral primary motor cortex (right: x = 60, y = 03, z = 18, non-parametric p_cluster_ FWE = 0.045; left: x = -57, y = -6, z = 24, non-parametric p_cluster_ = 0.0339).

**Figure 4.**
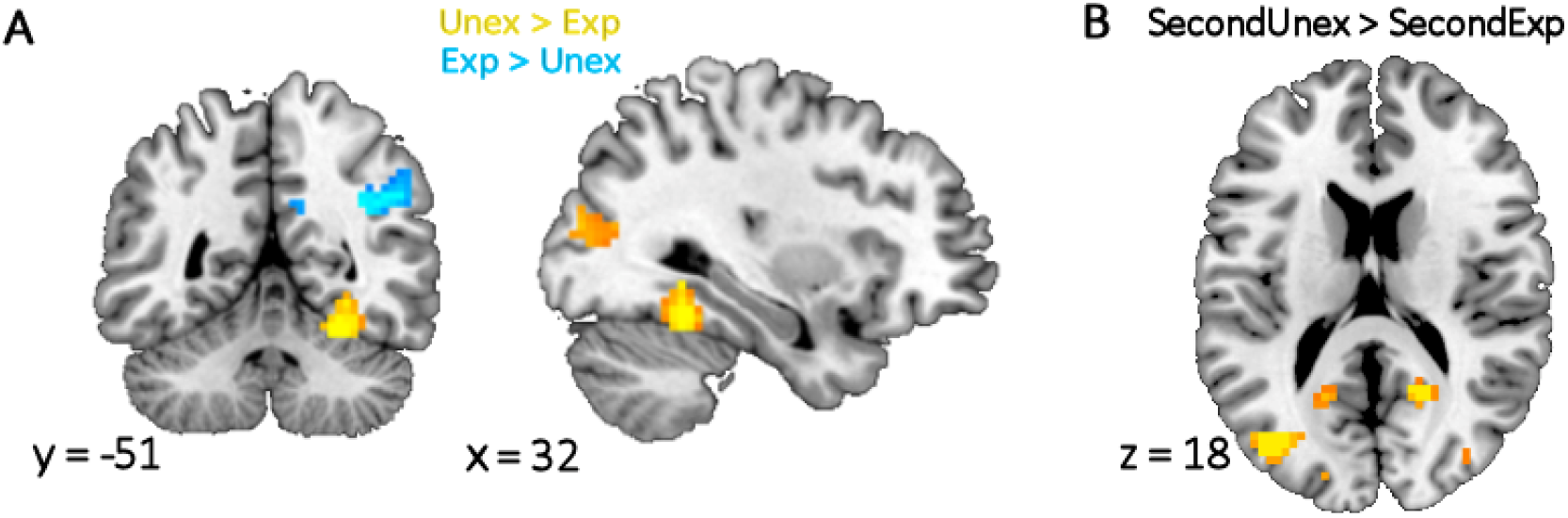
Overall fMRI expectation effects. A) Main effect of expectation. Unexpected events, compared to expected ones, engaged more activation in right middle occipital cortex and fusiform. Conversely, excepted events, compared to unexpected ones, engaged activation in right angular and supramarginal gyri. **B) Expectation by presentation order interaction**. Unexpected events presented second in the set engaged regions along the ventral visual stream. No effects were observed for first or third set events.

##### Occipital activation supports the interaction between expectation and memory performance for similar events

To unpack the overall unexpected > expected effect, we tested how contextual expectation interacted with successful recognition decisions (hits and CR), across presentation order. Whilst no differential neural responses were found for expected and unexpected hits or correct rejections of F2 events, we observed increased activation in the right inferior occipital gyrus (BA 19, x = 24, y = -81, z = -6, non-parametric p_cluster_ = 0.0345) for unexpected CR1 > expected CR1.

##### Increased perceptual load interacts with expectation status to engage ventral visual stream regions

Finally, we examined interactions between expectation status and presentation order (across set events; see Figure 4B). We observed again increased activity in bilateral occipital cortex (BA 19, x = -39, y = -78, z = 18, non-parametric p_cluster_ = 0.008, and BA 18, x = 18, y =-57, z = 21, non-parametric p_cluster_ = 0.009; x = 15, y = -87, z = -3, non-parametric p_cluster_ = 0.0129) for unexpected > expected events presented second in the set (no unexpected > expected effects were found for first or third set events).

## Discussion

The experience of surprise, or expectation violation, has a beneficial effect on learning, but whether surprise also triggers an encoding response even when the dominant goal is to retrieve, has remained unclear. In two experiments, we used a contextual expectation manipulation to better understand the dynamic nature of the adaptive memory mechanism triggered by expectation violation during retrieval, and its potential mnemonic consequences on hippocampal-dependent memory. We found that encountering unexpected events at retrieval, elicited increased involvement of regions along the ventral visual stream, even when memory performance was poor (U_prev_U_curr_). Interestingly, we also found a later beneficial effect of contextual surprise, such that the presentation of an unexpected event did not support its own recognition, but it did boost the correct recognition of the following, expected, and similar set events (U_prev_E_curr_). This behavioural effect was associated with increased activity in the hippocampus, midbrain dopaminergic regions (SN/VTA), and occipital cortex. Expected events, conversely, were associated with activity in retrieval-driven network regions. Given our replicated finding of the modulation of memory by previous unexpected events, the increased involvement of ventral visual stream regions, and previous research on expectation-modulated encoding, we postulate that engaging with unexpected information at retrieval engages an implicit bottom-up encoding mechanism (Figure 5). The consequences of this engagement become clear in the subsequent recognition trial, with a divergence in performance, and differential pattern of fMRI activation depending on whether the subsequent event was expected or unexpected.

**Figure 5.**
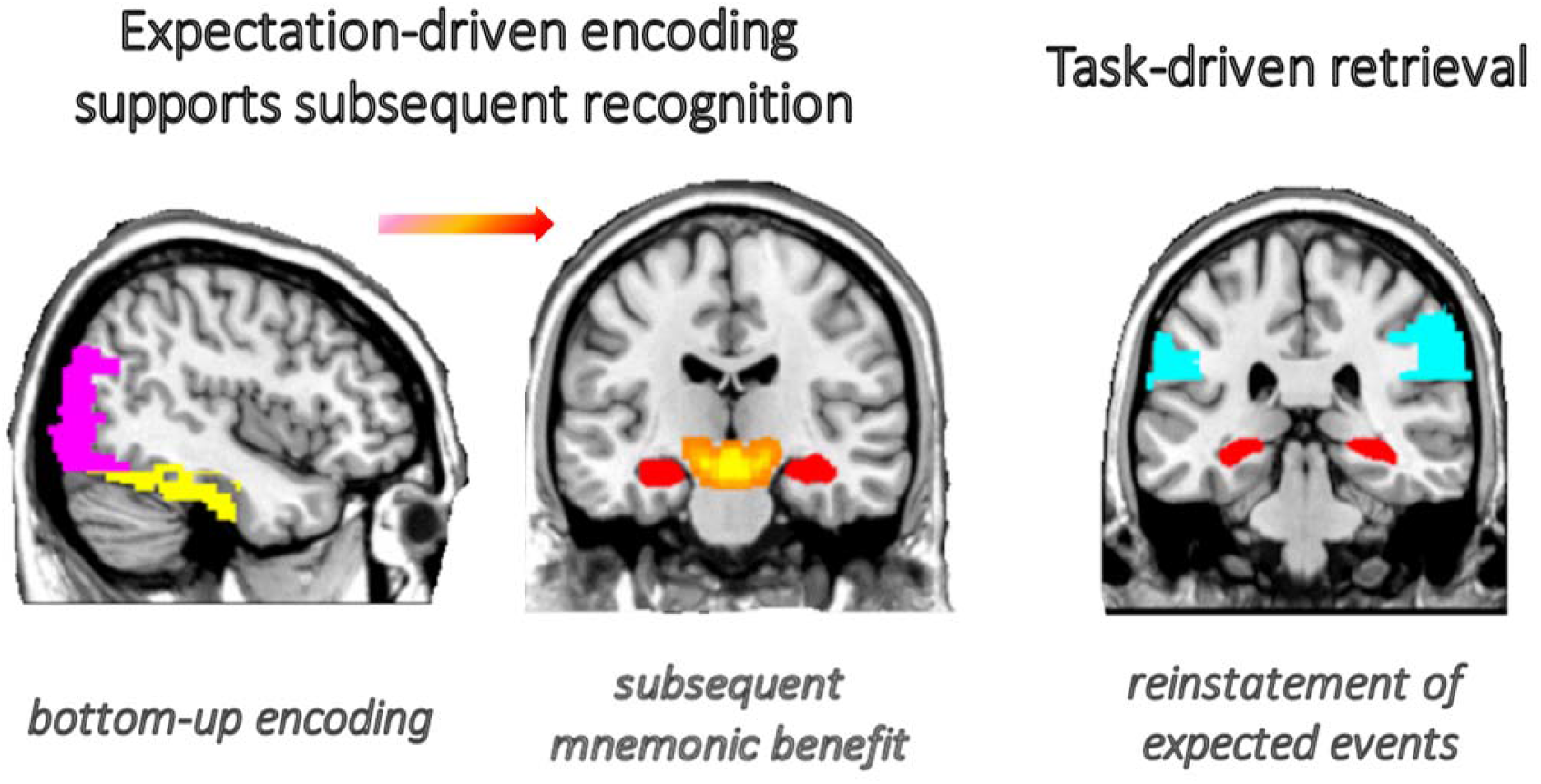
Illustration of the brain networks involved in processing unexpected and expected events. Left and middle: Expectation-driven, goal-irrelevant encoding. Expectation violation engages bottom-up processing along the ventral visual stream (inferior occipital in pink, fusiform in yellow), regardless of memory performance. The subsequent mnemonic consequences of this shift towards encoding involve the hippocampus (red) and midbrain (orange) dopaminergic regions, underlying subsequent beneficial memory performance. **Right: Task-driven retrieval**. In the absence of expectation violation (expected events), engagement of retrieval-driven network regions to support reinstatement and memory performance, in accordance with goal to retrieve.

Expectation violation is associated with improved memory performance, attributed to adaptive memory formation (Lisman and Grace, 2005; Kumaran and Maguire, 2007; Shohamy and Wagner, 2008), and impaired encoding of predictive information (Sherman and Turk-Browne, 2020). Our results support this view, but critically, extend it to account for retrieval effects. In line with the idea that increased weight is given to bottom-up inputs upon encountering a prediction error (Stoppel et al., 2009; Kafkas and Montaldi, 2018a), we found increased involvement of visual processing regions in occipital cortex and fusiform gyrus for unexpected events. These regions have been found to increase their activity with stronger levels of unexpected novelty (Kafkas and Montaldi, 2014), reflecting the increased perceptual processing of unexpected events. Although memory formation relies on bottom-up processing, evidence of an encoding mechanism requires that the mnemonic consequences of increased dependence on sensory inputs are demonstrated. Without subsequent mnemonic consequences, it could be argued that encountering an unexpected event only modulates online attention (Poort et al., 2022). Indeed, we observed an interaction whereby current memory performance was modulated by the previous unexpected occurrence of a similar event; when the previous event was expected, current expectation did not modulate performance (E_prev_E_curr_ ≈ E_prev_U_curr_), whereas when the previous event was unexpected, we found a divergence in performance (U_prev_E_curr_ > U_prev_U_curr_).

Taken together, these findings suggest that a surprise-driven increased weight on bottom-up inputs is goal-independent, but its mnemonic consequences appear to depend on the task at hand. During learning or exploration, further encoding supports later memory for the unexpected event (Li et al., 2003; Garrido et al., 2015; Long et al., 2016; Greve et al., 2017; Frank and Kafkas, 2021). When retrieval is the goal (as in the current paradigm), the implicit shift towards encoding, despite increased perceptual processing, results in numerically worse memory performance for the current to-be-retrieved information (Duncan et al., 2012; Kim et al., 2014). This is at odds with the notion that expectation violation always supports improved memory. Further support for the role played by perceptual load in engaging an encoding state, can be seen in the occipital and fusiform effects for those unexpected events presented second within the set sequence, compared to their expected counterparts. Recognition decisions for these events must overcome interference from the first set event, likely requiring increased perceptual processing to better compare the current sensory input with the stored representations.

Upon encountering the first unexpected event (U_prev_), a shift towards encoding, and away from retrieval, can explain why we do not observe a retrieval boost for these events. It is less obvious why this shift towards encoding produces better memory performance only for subsequent expected events. One possibility is that the initial expectation-violation driven shift towards encoding results in a sharper representation of the initial unexpected event (Gilboa and Moscovitch, 2021), optimising pattern completion of the second similar event (even when expected), as the delta similarity between the encoding and retrieval representations now stands out. Support for this account can be found in our fMRI findings; whilst occipital involvement was observed for both U_prev_E_curr_ and U_prev_U_curr_ events (i.e. independent of mnemonic consequence), only U_prev_E_curr_ events were associated with hippocampal and SN/VTA activation. This finding, together with the memory boost for U_prev_E_curr_ events, highlights the temporal contingency driven by U_prev_, as indexed by co-activation of SN/VTA and hippocampus (Kafkas and Montaldi, 2015). This co-activation is likely indicative of the expectation-driven (re)encoding of U_prev_ which then boosts memory for E_curr_.

Critically, interactions between current and previous events’ expectation were observed only for targets and F1 (i.e. the foil with most similarity to the encoded target). Moreover, these effects were unchanged by interfering events from the same set (F2, F3) or events from different sets presented during the task. That the expectation interactions are selective to high perceptual similarity, and are robust with respect to interference from other stimuli, suggests that a high perceptual and memorial load is required to trigger this encoding mechanism, consistent with, and extending, previous findings (Bein et al., 2020; Frank et al., 2020). In such situations, the ability to process and compare the fine details of current inputs and recently stored representations underpins correct recognition decisions (Yassa and Stark, 2011). Therefore, the triggering of enhanced perceptual processing by expectation violation serves an adaptive purpose (Stoppel et al., 2009; Hawco and Lepage, 2014). For less similar events, which are more readily recognised as new, a sharper representation, elicited by expectation violation, has little effect (Frank et al., 2020).

It is also important to consider how the shift towards encoding is manifested in the first event presentation; only first targets demonstrated a benefit for expected compared to unexpected previous events (and when examining only first set events). Whilst the increased hit rate for first expected targets is in line with our interpretation of the data, we did not observe a significant effect for F1 events. We suggest that this could be due to the intrinsic small differences in perceptual overlap between encountering a target and a very similar foil. It is possible that the increased difficulty associated with a first F1 event, driven by high but not full overlap with the encoded object, outweighs any potential effect of the implicit engagement of encoding. For targets, on the other hand there is a full perceptual overlap with the encoded object, that may facilitate recognition of expected targets, whereas expectation violation would deter it. Support for this interpretation can be found in the subsequent contrasts of the interaction in Experiment 1, where targets dominate the boost in U_prev_E_curr_, whereas F1 seem to drive the poor memory for U_prev_U_curr_. Furthermore, as discussed above, the lack of effects for lower-similarity foils suggests that perceptual load plays an important role in how expectation modulates memory processes. Given the robust behavioural interaction between subsequent events, and the complementary fMRI findings, we believe that an expectation-modulated shift towards encoding account best explains our data.

As the expectation manipulation took place at retrieval, it remains unclear whether encountering expected events resulted in task-relevant retrieval, or the active engagement of a retrieval state, irrespective of task demand. Although the engagement of temporo-parietal regions of a retrieval-driven network (Hayama et al., 2012) is indicative of reinstatement, this does not differentiate between the two alternatives. Future studies could orthogonalise expectation and memorial state, therefore allowing a factorial design of goal (encoding/retrieval) and expectation status. Examinations of shifts towards a retrieval state, perhaps coupled with designs optimised for functional and effective connectivity, will contribute to on-going efforts to explain how the hippocampus shifts between memory states (Colgin, 2016; Kafkas and Montaldi, 2018a; Bein et al., 2020). Whilst unexpected events during which you experience a high level of surprise are particularly memorable, it remains to be determined to what extent explicit awareness of surprise modulates this mechanism, and how memory demands might direct activity in the visual system.

In conclusion, we report novel evidence for the ubiquity of the adaptive encoding mechanism, here triggered at retrieval by expectation violation, resulting in differential effects on recognition performance. We propose that the increased demand on bottom-up occipital inputs, together with hippocampal-midbrain activations, are markers of an encoding state triggered by expectation violation, even in the absence of explicit reward or instruction. The complex temporal dynamics of the effects of this mechanism on memory demonstrate that the expectation-driven shift towards an encoding state engages increased perceptual processing, exerting a beneficial effect on correct recognition of subsequent similar events. These findings have important implications for our understanding of how our processing of sequential events, expected or unexpected, is modulated by the temporal dynamics of the event sequence.

## References

berg KC, Kramer EE, Schwartz S (2020) Interplay between midbrain and dorsal anterior cingulate regions arbitrates lingering reward effects on memory encoding. Nat Commun 11:1829 Available at: http://www.nature.com/articles/s41467-020-5542-z.

Aly M, Turk-Browne NB (2017) How Hippocampal Memory Shapes, and Is Shaped by, Attention. In: The Hippocampus from Cells to Systems, pp 369–403. Cham: Springer International Publishing. Available at: http://link.springer.com/10.1007/978-3-319-50406-3_12.

Ashburner J (2007) A fast diffeomorphic image registration algorithm. Neuroimage 38:95–113 Available at: https://linkinghub.elsevier.com/retrieve/pii/S1053811907005848.

Axmacher N, Cohen MX, Fell J, Haupt S, Dümpelmann M, Elger CE, Schlaepfer TE, Lenartz D, Sturm V, Ranganath C (2010) Intracranial EEG Correlates of Expectancy and Memory Formation in the Human Hippocampus and Nucleus Accumbens. Neuron 65:541–549.

Bates D, Mächler M, Bolker B, Walker S (2015) Fitting Linear Mixed-Effects Models Using lme4. J Stat Softw 67 Available at: http://arxiv.org/abs/1406.5823.

Bein O, Duncan K, Davachi L (2020) Mnemonic prediction errors bias hippocampal states. Nat Commun 11:3451 Available at: http://www.nature.com/articles/s41467-020-17287-1.

Buzsáki G (2002) Theta Oscillations in the Hippocampus. Neuron 33:325–340 Available at: http://linkinghub.elsevier.com/retrieve/pii/S089662730200586X.

Colgin LL (2016) Rhythms of the hippocampal network. Nat Rev Neurosci 17:239–249 Available at: http://www.nature.com/articles/nrn.2016.21.

Desikan RS, Ségonne F, Fischl B, Quinn BT, Dickerson BC, Blacker D, Buckner RL, Dale AM, Maguire RP, Hyman BT, Albert MS, Killiany RJ (2006) An automated labeling system for subdividing the human cerebral cortex on MRI scans into gyral based regions of interest. Neuroimage 31:968–980 Available at: https://linkinghub.elsevier.com/retrieve/pii/S1053811906000437.

Dimsdale-Zucker HR, Ranganath C (2019) Representational Similarity Analyses: A Practical Guide for Functional MRI Applications. Handb Behav Neurosci 28:509–525.

Duncan K, Sadanand A, Davachi L (2012) Memory’s Penumbra: Episodic Memory Decisions Induce Lingering Mnemonic Biases. Science (80-) 337:485–487.

Dunsmoor JE, Murty VP, Davachi L, Phelps EA (2015) Emotional learning selectively and retroactively strengthens memories for related events. Nature 520:345–348 Available at: http://www.nature.com/articles/nature14106.

Elston TW, Bilkey DK (2017) Anterior Cingulate Cortex Modulation of the Ventral Tegmental Area in an Effort Task. Cell Rep 19:2220–2230 Available at: https://linkinghub.elsevier.com/retrieve/pii/S2211124717307192.

Fox J (2003) Effect Displays in R for Generalised Linear Models. J Stat Softw 8 Available at: http://www.jstatsoft.org/v08/i15/.

Fox J, Weisberg S (2018) An R Companion to Applied Regression. Sage.

Frank D, Gray O, Montaldi D (2019) SOLID-Similar object and lure image database. Behav Res Methods Available at: http://link.springer.com/10.3758/s13428-019-01211-7.

Frank D, Kafkas A (2021) Expectation-driven novelty effects in episodic memory. Neurobiol Learn Mem 183:107466 Available at: https://linkinghub.elsevier.com/retrieve/pii/S1074742721000885.

Frank D, Montemurro MA, Montaldi D (2020) Pattern Separation Underpins Expectation-Modulated Memory. J Neurosci 40:3455–3464 Available at: http://www.jneurosci.org/lookup/doi/10.1523/JNEUROSCI.2047-19.2020.

Garrido MI, Barnes GR, Kumaran D, Maguire EA, Dolan RJ (2015) NeuroImage Ventromedial prefrontal cortex drives hippocampal theta oscillations induced by mismatch computations. Neuroimage 120:362–370 Available at: http://dx.doi.org/10.1016/j.neuroimage.2015.07.016.

Gilboa A, Moscovitch M (2021) No consolidation without representation: Correspondence between neural and psychological representations in recent and remote memory. Neuron 109:2239–2255 Available at: https://linkinghub.elsevier.com/retrieve/pii/S0896627321002919.

Greve A, Cooper E, Kaula A, Anderson MC, Henson RN (2017) Does prediction error drive one-shot declarative learning? J Mem Lang 94:149–165 Available at: http://dx.doi.org/10.1016/j.jml.2016.11.001.

Hasselmo ME, Bodelon C, Wyble BP (2002) A Proposed Function for Hippocampal Theta RhythmlZ: Separate Phases of Encoding and Retrieval Enhance Reversal of Prior Learning. Neural Comput 14:793–817.

Hawco C, Lepage M (2014) Overlapping patterns of neural activity for different forms of novelty in fMRI. Front Hum Neurosci 8 Available at: http://journal.frontiersin.org/article/10.3389/fnhum.2014.00699/abstract.

Hayama HR, Vilberg KL, Rugg MD (2012) Overlap between the Neural Correlates of Cued Recall and Source Memory: Evidence for a Generic Recollection Network? J Cogn Neurosci 24:1127–1137 Available at: http://www.mitpressjournals.org/doi/10.1162/jocn_a_00202.

Kafkas A (2021) Encoding-linked pupil response is modulated by expected and unexpected novelty: Implications for memory formation and neurotransmission. Neurobiol Learn Mem.

Kafkas A, Montaldi D (2014) Two separate, but interacting, neural systems for familiarity and novelty detection: A dual-route mechanism. Hippocampus 24:516–527 Available at: http://doi.wiley.com/10.1002/hipo.22241.

Kafkas A, Montaldi D (2015) Striatal and midbrain connectivity with the hippocampus selectively boosts memory for contextual novelty. Hippocampus 25:1262–1273 Available at: http://www.ncbi.nlm.nih.gov/pubmed/25708843.

Kafkas A, Montaldi D (2018a) How do memory systems detect and respond to novelty? Neurosci Lett 680:60–68 Available at: http://dx.doi.org/10.1016/j.neulet.2018.01.053.

Kafkas A, Montaldi D (2018b) Expectation affects learning and modulates memory experience at retrieval. Cognition 180:123–134 Available at: https://linkinghub.elsevier.com/retrieve/pii/S0010027718301938.

Kim G, Lewis-Peacock J a, Norman KA, Turk-Browne NB (2014) Pruning of memories by context-based prediction error. Proc Natl Acad Sci 111:8997–9002 Available at: http://www.pnas.org/cgi/doi/10.1073/pnas.1319438111.

Kumaran D, Maguire EA (2007) Which computational mechanisms operate in the hippocampus during novelty detection? Hippocampus 17:735–748 Available at: http://onlinelibrary.wiley.com/doi/10.1002/hipo.20207/abstract%5Cnpapers3://publication/doi/10.1002/hipo.20207.

Li S, Cullen WK, Anwyl R, Rowan MJ (2003) Dopamine-dependent facilitation of LTP induction in hippocampal CA1 by exposure to spatial novelty. Nat Neurosci 6:526–531.

Lisman JE, Grace A a. (2005) The Hippocampal-VTA Loop: Controlling the Entry of Information into Long-Term Memory. Neuron 46:703–713 Available at: http://linkinghub.elsevier.com/retrieve/pii/S0896627305003971.

Long NM, Lee H, Kuhl B a. (2016) Hippocampal Mismatch Signals Are Modulated by the Strength of Neural Predictions and Their Similarity to Outcomes. J Neurosci 36:12677–12687 Available at: http://www.jneurosci.org/lookup/doi/10.1523/JNEUROSCI.1850-16.2016.

Misaki M, Kim Y, Bandettini PA, Kriegeskorte N (2010) Comparison of multivariate classifiers and response normalizations for pattern-information fMRI. Neuroimage 53:103–118 Available at: https://linkinghub.elsevier.com/retrieve/pii/S1053811910007834.

Murty VP, Shermohammed M, Smith D V., Carter RM, Huettel SA, Adcock RA (2014) Resting state networks distinguish human ventral tegmental area from substantia nigra. Neuroimage 100:580–589 Available at: https://linkinghub.elsevier.com/retrieve/pii/S1053811914005242.

Oosterhof NN, Connolly AC, Haxby J V. (2016) CoSMoMVPA: Multi-Modal Multivariate Pattern Analysis of Neuroimaging Data in Matlab/GNU Octave. Front Neuroinform 10:1–27 Available at: http://journal.frontiersin.org/Article/10.3389/fninf.2016.00027/abstract.

Oyarzún JP, Packard PA, de Diego-Balaguer R, Fuentemilla L (2016) Motivated encoding selectively promotes memory for future inconsequential semantically-related events. Neurobiol Learn Mem 133:1–6.

Peirce JW (2007) PsychoPy-Psychophysics software in Python. J Neurosci Methods 162:8–13.

R Development Core Team (2008) R: A language and environment for statistical computing. Available at: http://www.r-project.org.

Richter FR, Chanales AJH, Kuhl BA (2016) Predicting the integration of overlapping memories by decoding mnemonic processing states during learning. Neuroimage 124:323–335 Available at: http://linkinghub.elsevier.com/retrieve/pii/S1053811915007703.

Searle SR, Speed FM, Milliken GA (1980) Population Marginal Means in the Linear Model: An Alternative to Least Squares Means. Am Stat 34:216–221 Available at: http://www.tandfonline.com/doi/abs/10.1080/00031305.1980.10483031.

Sherman BE, Turk-Browne NB (2020) Statistical prediction of the future impairs episodic encoding of the present. Proc Natl Acad Sci 117:22760–22770 Available at: http://www.pnas.org/lookup/doi/10.1073/pnas.2013291117.

Shohamy D, Adcock RA (2010) Dopamine and adaptive memory. Trends Cogn Sci 14:464–472.

Shohamy D, Wagner AD (2008) Integrating Memories in the Human Brain: Hippocampal-Midbrain Encoding of Overlapping Events. Neuron 60:378–389 Available at: http://dx.doi.org/10.1016/j.neuron.2008.09.023.

Stoppel CM, Boehler CN, Strumpf H, Heinze H-J, Hopf JM, Düzel E, Schoenfeld MA (2009) Neural correlates of exemplar novelty processing under different spatial attention conditions. Hum Brain Mapp 30:3759–3771 Available at: http://doi.wiley.com/10.1002/hbm.20804.

West BT, Welch KB, Galecki AT (2014) Linear Mixed Models. Chapman and Hall/CRC. Available at: https://www.taylorfrancis.com/books/9781466561021.

Wickham H (2009) ggplot2. New York, NY: Springer New York. Available at: http://link.springer.com/10.1007/978-0-387-98141-3.

Yassa MA, Stark CEL (2011) Pattern separation in the hippocampus. Trends Neurosci 34:515–525 Available at: http://linkinghub.elsevier.com/retrieve/pii/S0166223611001020.

